# Wireless electrocochleography in awake chinchillas: a model to study multimodal modulations at the peripheral level

**DOI:** 10.1101/2023.10.04.560827

**Authors:** Catherine Pérez-Valenzuela, Sergio Vicencio-Jiménez, Mia Memmel, Paul H. Delano, Diego Elgueda

## Abstract

The discovery and development of electrocochleography (ECochG) in animal models has been fundamental for its implementation in clinical audiology and neurotology. In our laboratory, the use of round-window ECochG recordings in chinchillas has allowed a better understanding of auditory efferent functioning. In previous works, we gave evidence of the corticofugal modulation of auditory-nerve and cochlear responses during visual attention and working memory. However, whether these cognitive top-down mechanisms to the most peripheral structures of the auditory pathway are also active during audiovisual crossmodal stimulation is unknown. Here, we introduce a new technique, wireless ECochG to record compound-action potentials of the auditory nerve (CAP), cochlear microphonics (CM), and round-window noise (RWN) in awake chinchillas during a paradigm of crossmodal (visual and auditory) stimulation. A total of four chinchillas were successfully recorded during the experimental protocol. There were non-significant differences in CAP, CM, and RWN amplitudes in response to auditory stimulation alone (clicks and tones) as compared to audio-visual crossmodal stimulation. These results suggest that cognition, such as attention or working memory, is needed for the corticofugal modulation of auditory-nerve and cochlear responses. In addition, we introduce the use of wireless ECochG in animal models as a useful tool for translational research.

## Introduction

In 1930, for the first time, bioelectrical potentials were recorded from the inner ear of cats (Wever and Bray, 1930). In this seminal article, Wever and Bray described an alternating potential that allowed them to communicate from inside to outside a soundproof room as if it were a microphone. Nowadays, we know it corresponds to the cochlear microphonic potential (CM) (Eggermont, 2017). Since this first electrocochleography (ECochG) report, an increasing number of studies in animal models have allowed the understanding of the origin of these extracellular potentials (Dallos and Cheatham, 1976; Siegel, 1986).

It is widely accepted that the bioelectric signals recorded with ECochG are originated from cochlear and neural sources, including cochlear microphonics from cochlear hair cells (Wever and Bray, 1930), summating potentials from cochlear ramps (Dallos et al, 1970), compound action potentials (CAP) of the auditory nerve (Davis et al., 1952), neurophonic potentials from auditory nerve and brainstem (Snyder and Schreiner, 1984), and the ensemble background activity (also known as round window noise (RWN)) which reflects spontaneous neural activity (Dolan et al., 1990; Searchfield et al., 2004). This variety of responses can be isolated using different recording techniques and analytic methods, such as alternating polarity stimuli or different signal filtering (Ferraro and Krishnan, 1997), and have been used for research in several fields in auditory neuroscience (Galambos, 1956; Elgueda et al., 2011; Dragicevic et al., 2015), and to propose multiple applications for clinical practice in humans (Eggermont, 2017; Gibson, 2017).

### Use of electrocochleography to understand auditory efferent function

The auditory efferent system is a neural network composed of several feedback loops that connect the central nervous system with auditory-nerve neurons and cochlear hair cells (Terreros and Delano, 2015; Elgueda and Delano, 2020). The final connections between the brainstem and auditory peripheral levels are made by olivocochlear neurons (Warr and Guinan, 1979), including the medial and lateral olivocochlear synapses with outer hair cells and auditory-nerve neurons, correspondingly (Lauer et al., 2022). The possible functions of the olivocochlear system and its role in audition have been a matter of study for several decades, including a number of studies that used animal models and electrocochleographic potentials, mainly CAP and CM responses (Galambos, 1956; Smith et al., 1994; Vetter et al., 1999; Elgueda et al., 2011; Verschooten et al., 2012; Aedo et al., 2015).

Cochlear sensitivity modulation through selective attention has been raised as a possible function of the auditory efferent system, a hypothesis supported by a series of studies performed in animal models, including cats (Hernandez-Peón et al., 1956; Oatman 1971), chinchillas (Delano et al., 2007; Bowen et al., 2020; Vicencio-Jimenez et al., 2021), and mice (Terreros et al., 2016). For instance, in Delano et al., (2007) we showed that during periods of visual selective attention, CAP responses were reduced, while CM responses were augmented. This paradoxical effect in CAP and CM responses is considered a signature of medial olivocochlear activation (Galambos, 1956; Fex 1959; Robertson 2009; Elgueda et al., 2011; Jennings and Aviles 2023). Altogether animal studies gave evidence that the corticofugal projections of the auditory efferent system can modulate afferent responses during crossmodal attention paradigms, in which animals focus their attention on visual stimuli while suppressing responses to auditory distractors.

Here, we hypothesized that crossmodal modulations on auditory-nerve and cochlear microphonic responses by auditory efferents are also evident in non-attending awake animals. To answer this proposal, we used wireless ECochG to record CM, CAP, and RWN potentials in awake chinchillas during crossmodal stimulation with auditory and visual stimuli.

## Methods

### Animals

Four adult chinchillas (*Chinchilla lanigera*, 4 ± 1 years of age) weighing 400-700 g were used in this study. Chinchillas were individually housed in temperature and humidity-controlled cages located inside a room where the dark/light cycle was inverted (lights on from 8 P.M. to 8 A.M.). Throughout the experimental phase, they were granted unrestricted access to water while being subjected to food deprivation, with their weight maintained at 85 to 90% of their free-feeding weight. Animals spent at least 3 hours per week in an enrichment environment, where they could exercise and bathe. All experimental procedures were approved by the local institutional animal care and use bioethical committee (Comite de Bioetica Animal, permit number #0844, Facultad de Medicina, Universidad de Chile) and adhered strictly to the Guidelines for the Care and Use of Laboratory Animals (National Research Council, 2011).

### Surgery

The weight of the animals was monitored daily, and only chinchillas whose weight variation did not exceed 10% during the week prior to surgery qualified for the procedure. As a preoperative protocol, the animals received a dose of ketoprofen 5 mg/kg (i.m.) and atropine 0.4 mg/kg (i.m.) 30 min before surgery. Chinchillas were anesthetized with ketamine (20 mg/kg, i.m.) and xylazine (3 mg/kg, i.m.). Ophthalmic chloramphenicol cream was applied to the eyes to prevent corneal injury. The level of anesthesia was maintained by giving repeated half doses every 45-60 minutes or as needed, as judged by the foot withdrawal reflex. Rectal temperature was maintained at 35-37°C using a heating pad. Chinchillas were secured in a stereotaxic apparatus, where electrodes were implanted bilaterally in the round window of each cochlea and unilaterally in the left temporal cortex (cortical data not shown in this article). A small 1 mm dorsal opening was made on the auditory bullae for round window electrode placement and low-impedance, 100 μm nichrome electrodes were inserted into the cochlear round window of each cochlea, and connected to an omnetics connector placed on the skull. Cyanoacrylate and dental acrylic cement were used to secure the electrode and connector to the skull. Postoperatively, the animals were supplemented with a semi-liquid diet consisting of pureed fruits and vegetables and were also given 2.5 mg/kg of oral ketoprofen for three days. Chinchillas were allowed to recover for seven days before wireless recordings.

### In vivo wireless electrophysiology

All recordings were acquired from awake, freely moving animals utilizing a wireless system (W2100, Multichannel Systems MCS GmbH, Harvard Bioscience, Inc., Reutlingen, Germany). The system uses a 16-channel headstage that was connected to the omnetics connector during the behavioral paradigm (Fig. 1). The wireless system has an amplifier bandwidth ranging from 1 Hz to 5 kHz, and a sampling rate of 20 kHz (with a gain of 101X, 1 GΩ input impedance, 16-bit resolution, ± 12.4 mV input voltage range, an input noise below <1.9 μVRMS, and 5 m of wireless link distance). The Multichannel Suite software was used for data acquisition. Electrodes exhibiting poor signal quality and motion artifacts were visually identified offline and discarded from analyses.

**Fig. 1.**
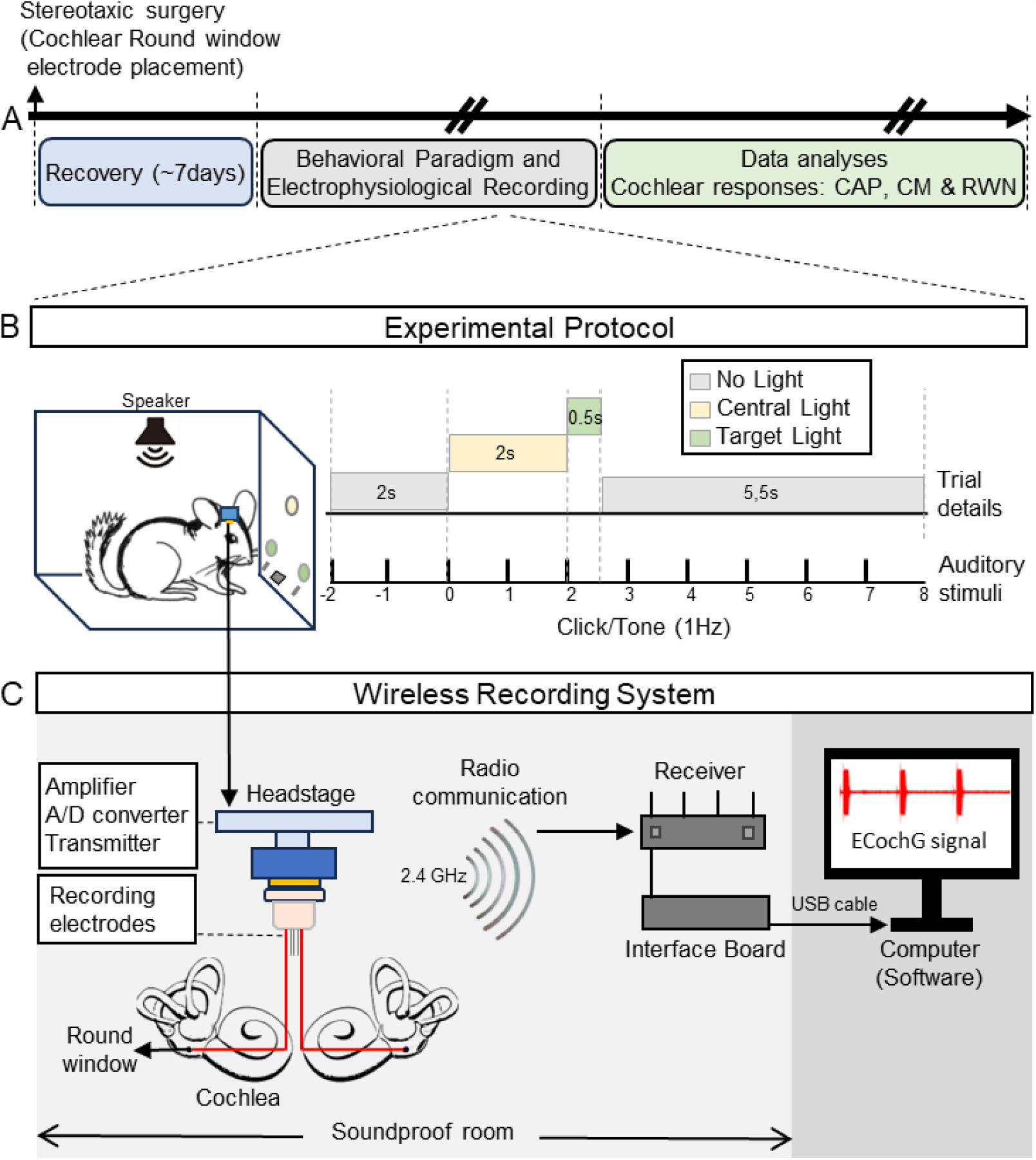
Experimental protocol. **(A)** Timeline of the experimental procedures, a chronic surgery for implanting round-window electrodes. **(B)** Behavioral conditions during awake recordings. Chinchillas were placed in an operant mesh box where they could move freely. Sound stimuli consisting of a click and a tone were presented at a 1 Hz rate. Statistical analyses were performed comparing the first two seconds (no light) with the two seconds with the neutral cue light in omitted trials. **(C)** Schematic diagram of wireless ECochG recordings. The electrodes were connected with the head stage only for recordings. The head stage contains an amplifier and A/D converter and transmitter that sends the ECochG signal at 20 kHz/s at 2.4 GHz. The receiver is located inside the soundproof room and is connected to the outside computer via a USB cable.

### Behavioral paradigm

All animals underwent training for 3 to 4 months, 5 days per week, in a two-choice visual discrimination task using an operant cage inside a dark double-walled sound-attenuating room (Delano et al., 2007). The front panel of the cage was equipped with three lights: a central light serving as a neutral cue located above a food dispenser and two lateral lights functioning as targets, positioned above the right and left levers (Delano et al., 2007).

The behavioral paradigm was controlled with custom software written in C language (LabWindows/CVI, National Instruments, Austin TX, USA) and a National Instruments DAQ board (National Instruments, PCI-6040E) housed in a desktop computer. Each trial followed a specific sequence: it initiated with 2 seconds of darkness (no light), followed by a 2-second warning period marked by the activation of a neutral cue (central light). Subsequently, one of the target lights (either the right or left, selected randomly) turned on for a period 0.5 seconds. The animals had 5.5 seconds to respond by pressing a lever under the target light in order to receive a food pellet reward. Simultaneously to the visual stimulation, background sound stimuli were presented, consisting of a continuous presentation of a click (100 microseconds) followed (50 ms) by a 2 kHz tone (100 ms duration), presented at a 1 Hz presentation rate. These sounds were presented with phase alternation in odd and even trials to allow for CAP and CM separate averaging procedures.

Chinchillas were trained to press one of the two levers, corresponding to each target light (left/right lever). Correct responses were rewarded with a food pellet, while incorrect responses were punished with a 40 s time-out in darkness. The trials in which the chinchillas did not respond were defined as omissions and were selected to analyze cochlear responses in an inattentive state with and without visual stimulation presented in this article (Fig. 1).

### Data Analyses

ECochG signals were acquired at 20 kHz and analyzed offline with scripts written in MATLAB (R2021b, Mathworks, Natick, MA, USA). Signals were filtered using a forward-reverse 4th-order Butterworth band-pass filter (100 -5000 Hz). CAP signal was measured from click responses as the peak-to-peak amplitude (μV) in a time window of 2-7 ms after click onset. CM amplitude was analyzed from tone responses, and computed from the power spectral density (PSD) of the signal, obtained with Welch’s periodogram method, and measuring the peak power at the tone stimulation frequency (2 kHz). RWN power was measured from the signal obtained during the silent periods (800 ms width) between sound stimuli (see Fig. 2) and integrating the PSD (using the trapezoidal method) between 800 and 1200 Hz.

**Fig. 2.**
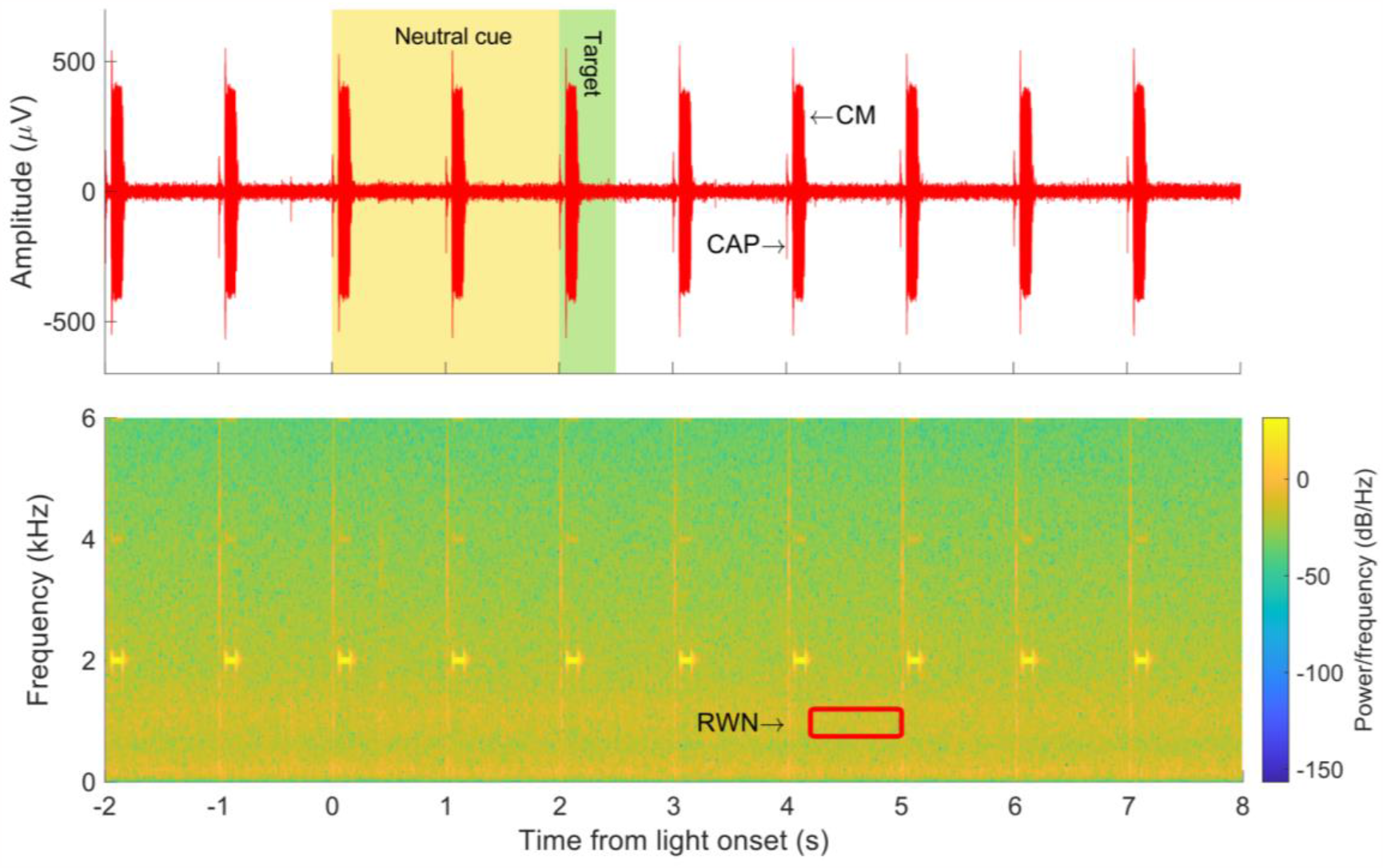
Example of single-trial wireless ECochG signal and its corresponding time spectrogram. The red trace in the upper panel shows CAP and CM responses to a 1 Hz-rate click followed by 2 kHz tone stimuli. A sound file saved from this signal can be found as Supplementary material 1. RWN responses were obtained from the spectrum (800-1200 Hz) of the silent time windows between sound presentations, one of them shown with a red rectangle.

CAP, CM, and RWN amplitudes were referenced in dB (20*log_10_[amplitude/reference amplitude]) relative to the mean amplitude obtained before the neutral cue onset (responses at -2 and -1 seconds before the onset of the neutral cue). Variations in CAP, CM, and RWN mean dB amplitudes measured during the neutral cue light (at 1 and 2 s relative to the central light onset (neutral cue)) were analyzed with a one-way repeated-measures analysis of variance (ANOVA). Because responses to the auditory stimuli presented at 0 ms were too fast for evaluating behavioral changes, the CAP, CM, and RWN responses at the light onset (0 s) were not considered for the statistical analyses. In the statistical analyses, a p-value < 0.05 was considered as significant.

## Results

Four chinchillas were successfully implanted for wireless ECochG and recorded in awake conditions in a crossmodal paradigm with auditory and visual stimulation during omitted trials. Figure 2 shows an example of the signal obtained in a single trial of the behavioral paradigm using wireless electrocochleography in one chinchilla, illustrating the quality of the recordings along the experimental protocol at the single trial level. A sound file corresponding to the signal shown in Fig. 2 is available as Supplementary Material 1. Fig. 3A shows the mean CAP and CM obtained from the ECochG signal over 25 trials and Fig. 3B displays the spectrum of the signal obtained from the period between sound presentations, which was used to compute RWN power.

**Fig. 3.**
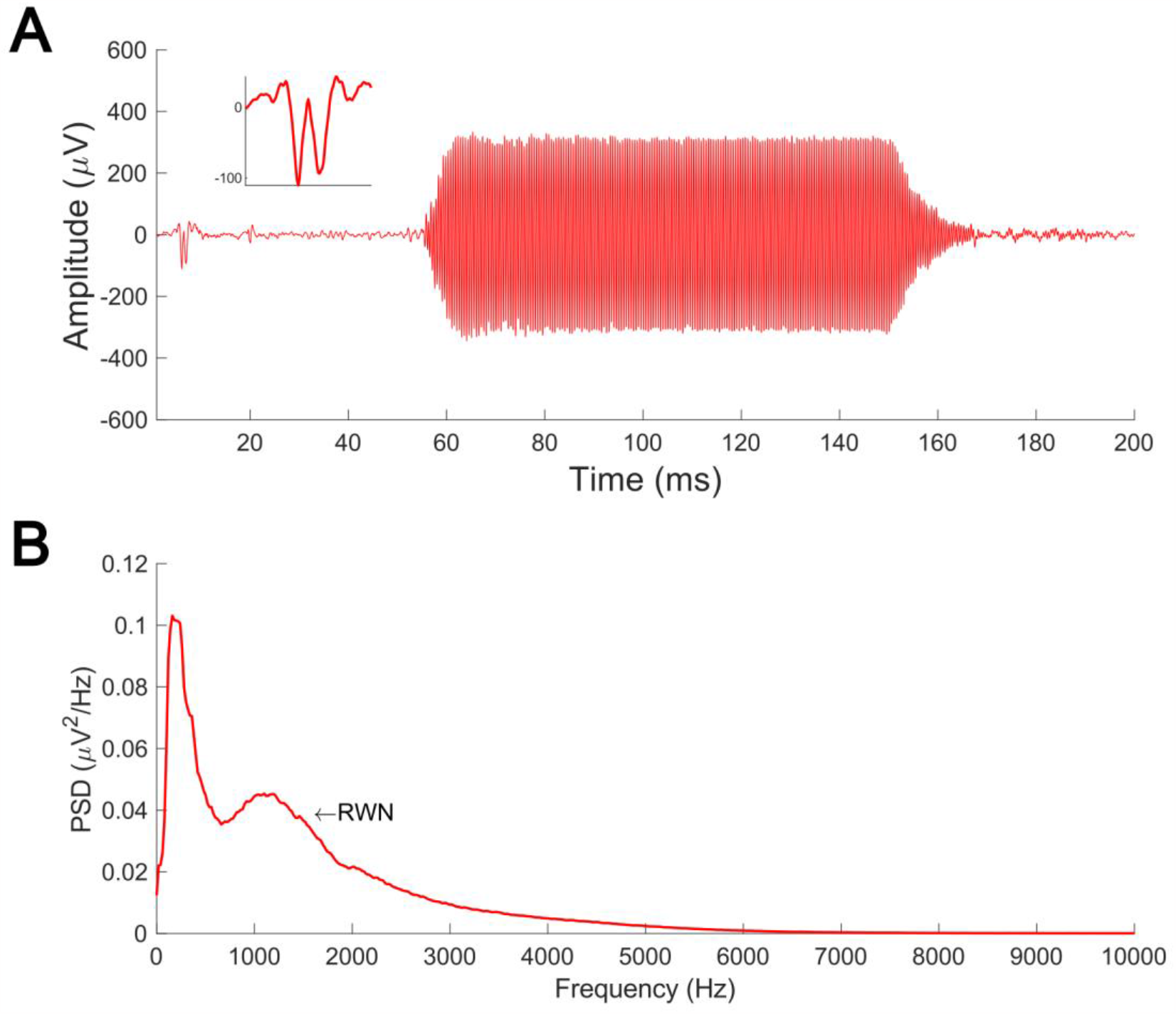
Wireless ECochG averaged responses. **(A):** Average CAP and CM responses to click and 2 kHz tone. These responses were averaged from 25 trials. The inset displays the CAP response. **(B):** RWN power spectral density (PSD). The magnitude of round window noise was computed by the integral value between 800 and 1200 Hz in the averaged spectrum.

CAP, CM, and RWN responses were averaged along omitted trials (i.e. trials where the animal did not respond) and referenced in dB to the baseline period (-2 to 0 seconds in the experimental protocol). The dB-referenced amplitudes were computed for the four chinchillas along time to compare ECochG responses in a total of 23 recordings (of 25 to 110 trials each). The repeated measures ANOVA analysis yielded non-significant differences comparing CAP, CM, and RWN responses before and during the neutral cue (CAP: F(3)=0.11476, p=0.95113; CM: F(3)=1.4998, p=0.22324; RWN: F(3)=1.4357, p = 0.24029). Fig. 4 shows boxplots of the grand average CAP, CM, and RWN amplitude data from the four animals along the experimental protocol.

**Fig. 4.**
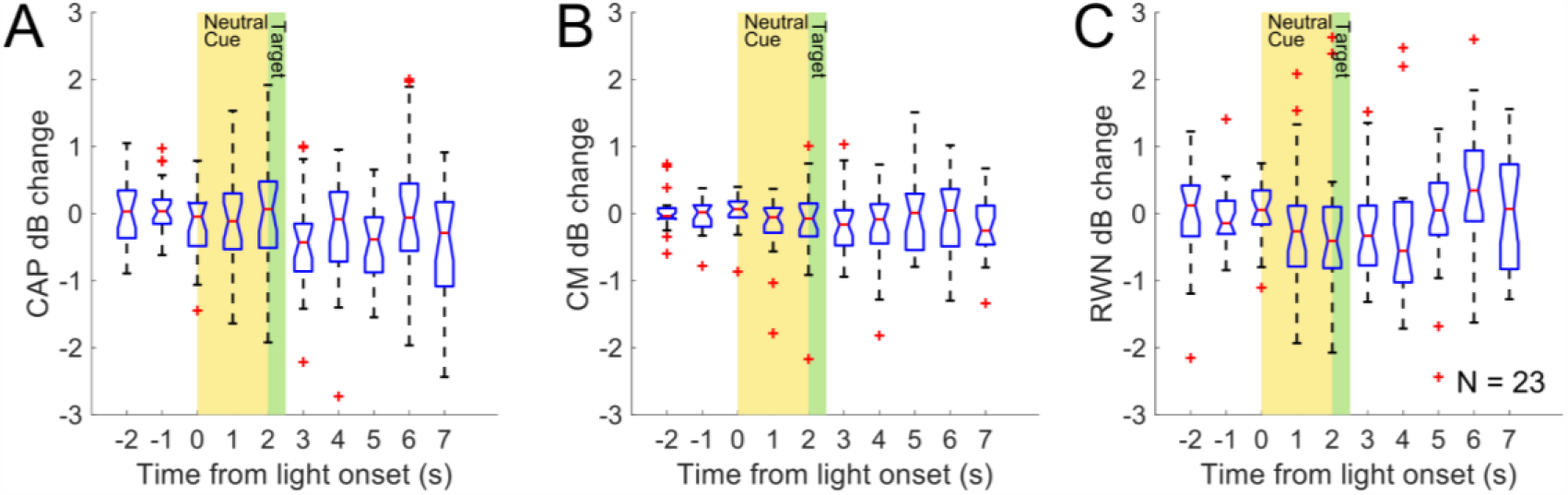
**(A)** CAP, **(B)** CM, and **(C)** RWN grand average amplitude changes during the experimental protocol. There were non-significant changes in the amplitudes of these ECochG potentials during crossmodal audio-visual stimulation (the period between 0 to 2 seconds (neutral cue period)) in omitted trials. Boxplots represent CAP, CM, and RWN amplitude changes referenced in dB to baseline levels (from -2 to 0 seconds), red crosses illustrate outliers.

## Discussion

In the present investigation, we introduced a wireless ECochG technique to record auditory-nerve CAPs, cochlear microphonics, and round window noise in awake chinchillas during a crossmodal (visual and auditory) stimulation paradigm (Fig. 1). This widens the possibility to study these physiological phenomena in chinchillas that are actively interacting with their environment. In particular, we have been especially interested in developing this model to further our understanding of auditory efferent physiology in chinchillas.

### Chinchillas as a model for auditory research

The chinchilla (*Chinchilla lanigera*) is a rodent from Chile and Argentina (Spotorno et al., 2004) widely used for auditory research due to several advantages as an animal model. Some of these include an auditory range similar to humans (Heffner and Heffner, 1991), relatively easy access to middle and inner ear surgery, and access to the cerebral cortex (Delano et al., 2007; Delano et al., 2010; Verschooten et al., 2012; León et al., 2012; Dragicevic et al., 2015), and the possibility to evaluate behavioral research (Bowen et al., 2020; Vicencio-Jimenez et al., 2021). In addition, they are social animals that emit vocalizations that can be used for ecological stimulation with natural stimuli (Moreno-Gómez et al., 2015; Bowen et al., 2020; Vicencio-Jimenez et al., 2021).

For this study, we were interested in exploring the potential impact of passive perception of visual stimuli on peripheral auditory activity, evaluated by CAP, CM, and RWN responses during omission trials. Our results showed that auditory stimulation alone, in the form of clicks and 2 kHz tones, elicited CAP, CM, and RWN potentials that were not significantly different in amplitude from those observed during crossmodal audio-visual stimulation (Fig. 4).

When placed in the context of previous research, the absence of an inattentive visual crossmodal influence on cochlear activity takes on special significance. It has been proposed that when animals focus their attention on a visual stimulus, the auditory efferent system suppresses afferent responses, possibly facilitating the direction of attentional resources to the relevant stimulus (Lauer et al., 2022). This selective attentional regulation of efferent activity is supported by research in cats (Hernández-Peón et al., 1956; Oatman 1971), chinchillas (Delano et al., 2007; Bowen et al., 2020; Vicencio-Jiménez et al., 2021) and mice (Terreros et al., 2016). In humans, the evidence is less consistent, although there are results showing a correlation between selective attention, working memory and modulation of cochlear activity (Wittekindt et al., 2014; Dragicevic et al., 2019; Marcenaro et al., 2021).

As mentioned above, our results show that under conditions of passive perception (when the animal is not actively attending to the stimulus during omission trials), these crossmodal effects are not evident. This underlines the specificity of cognitive processes on the efferent pathway modulation. In this context, the task we used in this research is particularly suitable for asserting that the chinchillas were not engaged with the visual stimulus. The animals were trained for a task in which visual stimuli are needed for obtaining a food reward. As we have shown previously, chinchillas are able to actively direct their attentional resources toward visual stimuli, ignoring auditory distractors (Delano et al., 2007, Bowen et al., 2020). For the purposes of this research, we were interested in observing only those events in which the chinchillas omitted their response, which allowed us to assume that in such trials they did not attend to visual information in pursuit of a goal. Thus, our results support the idea that it is not visual sensory stimulation (or other crossmodal stimuli) per se that modulate efferent auditory activity, showing that cognitive mechanisms are needed to activate corticofugal pathways to the cochlear receptor.

### Limitations

The present study has certain limitations that are important to consider when interpreting its results. One of the key limitations lies in the small number of animals used in the research, being limited to a group of only four chinchillas. Regarding this, it should also be noted that analysis of correct or incorrect responses was not included due to the low number of available trials. The animals analyzed in this study did not reach the level of performance necessary to include such data.

In addition, another aspect to consider is that the stimuli presented during the study consisted of highly predictable and non-natural stimuli. This predictability may have influenced subjects’ responses, limiting the study’s ability to assess how individuals would react in less predictable or more realistic situations.

### Future projections

One promising avenue would be to incorporate recordings from the auditory cortex or other regions of the central nervous system to analyze how neural activity changes during passive or active multimodal perception and how it correlates with auditory peripheral activity. This would allow further exploration of functional connectivity between different sensory systems and how they adapt in response to crossmodal stimuli. In addition, it could be especially valuable to use this recording technique to investigate active goal-directed responses in more naturalistic settings, which would involve, for instance, studying how the auditory cortex modulates the efferent system when the animal is attending to a stimulus from another sensory modality. Wireless ECochG could be obtained simultaneously with cortical recordings, which might allow the investigation of several questions in freely-moving animals engaged in natural behaviors.

## Conclusion

The present evidence supports the idea that corticofugal modulation of the olivocochlear system only occurs during active engagement with surrounding stimuli. These findings provide insights into the connection between cognitive processes, our active sensing of the environment, and the ability to regulate auditory nerve and cochlear responses. Also, wireless recordings have the potential to integrate cochlear and neural simultaneous recordings in freely moving animals engaged in natural settings, which may provide significant insights into the biological basis of behavior and perception.

## Acknowledgements

Funded by ANID, FONDECYT 11190278 to D.E.; ANID, FONDECYT 1220607 to P.H.D; and Fondo BASAL ANID FB0008 to P.H.D. We thank Fernando Vergara and Giuliana Bucci-Mansilla for their technical assistance.

## Notes

### Competing Interest Statement

The authors have declared no competing interest.

